# A High throughput method for egg size measurement in *Drosophila*

**DOI:** 10.1101/2022.09.13.507758

**Authors:** Neda Barghi, Claudia Ramirez-Lanzas

**Affiliations:** Institut für Populationsgenetik, Vetmeduni Vienna, Vienna, Austria; Vienna Graduate School of Population Genetics, Vetmeduni Vienna, Vienna, Austria

**Keywords:** Egg size, Large particle flow cytometry, *Drosophila*

## Abstract

Life-history traits are used as proxies of fitness in insects including *Drosophila*. Egg size is an adaptive and ecologically important trait potentially with genetic variation across different populations. However, the low throughput of manual measurement of egg size has hampered the widespread use of this trait in evolutionary biology and population genetics. We established a method for accurate and high throughput measurement of *Drosophila* egg size using large particle flow cytometry (LPFC). The size estimates using LPFC are accurate and highly correlated with the manual measurements. The measurement of egg size is high throughput (average of 214 eggs measured per minute) and viable eggs of a specific size can be sorted rapidly (average of 70 eggs per minute). Sorting by LPFC does not reduce the survival of eggs making it a suitable approach for sorting eggs for downstream analyses. This protocol can be applied to any organism within the detectable size range (10-1500 μm) of the large particle flow cytometers. We discuss the potential applications of this method and provide recommendations for optimizing the protocol for other organisms.

## Introduction

Egg size is an important trait in insects that have evolved in response to developmental and ecological pressures (Church et al., 2019). Egg size and other life-history traits (e.g. survival, longevity, and female fecundity) are proxies of fitness in insects. In *Drosophila melanogaster*, egg size affects other life-history traits e.g. embryonic viability, hatching rate, and embryonic development (Azevedo et al., 1997), and morphological features such as the embryonic anterior-posterior patterning (Huang et al., 2020; Lott et al., 2007). Larval crowding (Venkitachalam et al., 2022) and environmental factors such as temperature influence *D. melanogaster* egg size (Avelar, 1993; Azevedo et al., 1996; Imai, 1932). *D. melanogaster* egg size increases with latitude in Australia and South America (Azevedo et al., 1996) suggesting that it is an adaptive trait under stabilizing selection (Azevedo et al., 1997). *Drosophila* is one of the most widely used sexual multicellular organisms in evolutionary biology. This popularity is partly due to *Drosophila’s* short generation time and ease of maintenance in laboratory conditions. Despite the widespread use of *Drosophila*, egg size is rarely studied in evolutionary and population biology studies partly due to the low throughput of the manual measurement of egg size.

Traditionally, the size of eggs were measured when viewed under a microscope or stereoscope (Azevedo et al., 1996, 1997; Imai, 1932; Markow et al., 2009; Warren, 1924). Alternatively, the eggs are photographed under a microscope and their sizes are manually measured from the images (Huang et al., 2020; Jha et al., 2015; Lott et al., 2007; Miles et al., 2011). Recently, few image analysis tools are developed to automatically calculate the size of objects (Lürig, 2022; Waithe et al., 2015). The methods that rely on manual measurements are time-consuming and tedious because the eggs should be manually transferred and arranged in proper orientation before images are taken. Measuring the egg size manually is also error-prone due to the experimenter’s bias. The image analysis pipelines also require the specimens to be photographed against a background with specific color and brightness (Lürig, 2022). Moreover, manual measurements or image analysis tools usually cannot ensure the recovery of viable eggs that can be used for downstream analyses.

Flow cytometry allows fast and accurate sorting and size measurement of small objects up to 200 μm. But the average size of eggs in *D. melanogaster* species subgroup ranges from 400 to 600 μm (Imai, 1932; Lott et al., 2007; Markow et al., 2009; Venkitachalam et al., 2022; Warren, 1924) which is too large to be processed with the traditional flow cytometers. The advent of large particle flow cytometry (LPFC) has made the measurement of live objects up to 1500 μm possible. The LPFC systems operate at a slower flow rate and lower pressures compared to the traditional flow cytometers to avoid disruptive shear forces. LPFC has been used for the analysis of fluorescent-labeled or transgenic organisms e.g. egg of nematodes (Pillai & Dandurand, 2021), *D. melanogaster* (Furlong et al., 2001) and *Anopheles gambiae* (Marois et al., 2012), and coral larvae (Randall et al., 2020). Non-fluorescent labeled objects can also be analyzed and sorted on LPFC solely based on their size. However, in the absence of fluorescence labeling, the shape of objects affects the accuracy of size estimation. For example, the shape of *Arabidopsis* seeds is prolate spheroid or cardioid (Cervantes et al., 2010) thus the size measurement of *Arabidopsis* seeds is almost independent of the orientation of seeds (Morales et al., 2020). However, for objects with oblate or prolate ellipsoid shapes such as *Drosophila* eggs the estimated size may vary depending on the orientation of the objects in the flow cytometer. The size of the object is properly measured only when the objects are oriented along their axis but may be underestimated if the objects don’t pass along their axis (Fig. 1A-B).

**Fig. 1.**
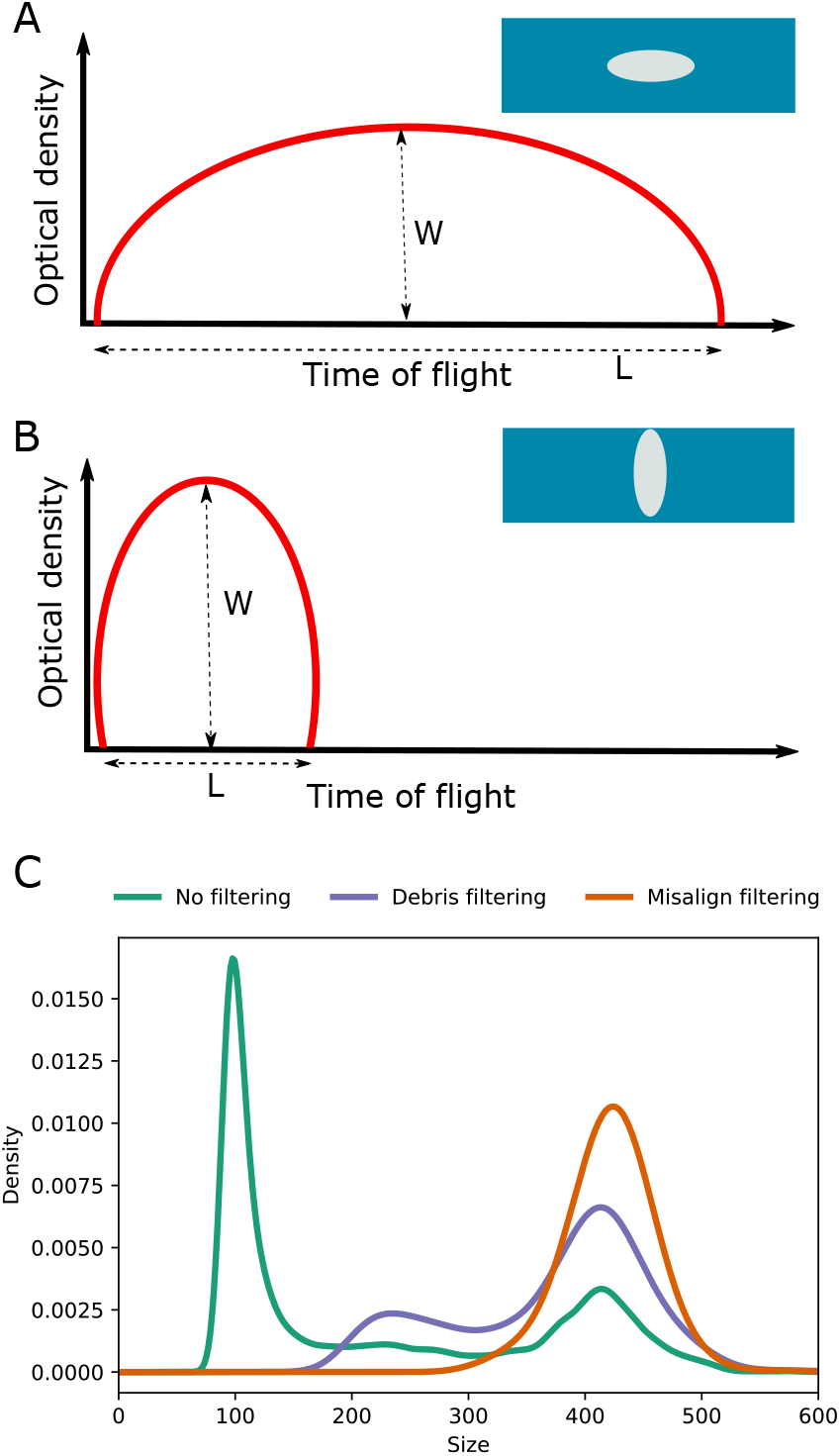
In silico filtering of debris and misaligned objects using the optical density profiles. A) *Drosophila* eggs are elliptical, and accurate size measurement based on time of flight (TOF) is obtained when an egg is aligned on its long axis. B) The size of a misaligned egg, i.e. aligned on its short axis, is underestimated. The misaligned eggs can be identified based on the ellipticalness index (EI) and W/L ratio where W is the maximum optical density of an object and L is the number of extinction measurements along an object’s time of flight. C) The initial size distribution is bimodal (No filtering). The 1^st^ mode corresponds to debris and yeast particles, and ‘Debris filtering’ remove these small objects. Further filtering steps based on EI and W/L parameters (Misalign filtering) removed the misaligned eggs. Size is in μm units. The egg length distribution of sample Dsim196 is shown in C: The statistics of egg length distribution is presented in Table S1 and S2, and the histogram of egg length distribution is depicted in Fig. S3.

In this study, we developed a method to measure the length of *Drosophila* eggs using LPFC. In this method, we filter out the eggs that are not oriented properly along their long axis in-silico and accurately measure the length distribution of *Drosophila* eggs. The measurement of egg length is accurate, fast, and has high throughput. Eggs of a specific size range can be sorted accurately and developed into adults with no reduction in viability. We discuss the potential of using this approach to investigate the natural variation and the genetic basis of egg size in *Drosophila*. LPFC can be used to measure the size of any organism within the detectable range (10-1500 μm) of this instrument. We provide recommendations for optimizing the use of LPFC for other organisms in particular when their shape deviates from spheroid.

## Materials and Methods

### Egg length as a proxy of egg size

Studies on *D. melanogaster* have used the egg length (Huang et al., 2020; Imai, 1932; Lott et al., 2007; Warren, 1924), egg volume (Avelar, 1993; Azevedo et al., 1996, 1997; Jha et al., 2015; Miles et al., 2011; Schwarzkopf et al., 1999) or both the length and width of egg (Markow et al., 2009) as proxies of size. Egg volume is a composite measure typically calculated using the formula 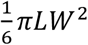 where *L*, length, corresponds to the axis of rotational symmetry and *W*, width, is the axis perpendicular to length (Church et al., 2019). Egg volume, together with the total number of eggs, can be used as a measure of reproductive investment (Avelar, 1993). However, when egg volume is used as a proxy of size, the inter- or intra-specific differences cannot be directly attributed to the variation in its components, i.e. length and width.

An advantage of measuring the length or width of eggs as size proxies is that they can be evaluated as distinct traits. These attributes of egg size may contribute differently to the variation in egg size across different species and populations. For example, in artificially selected populations for higher egg volume, eggs are shown to be longer (Miles et al., 2011). Functional validation of a few candidate genes identified in these populations has also shown that these genes affected egg length (Jha et al., 2015). This suggests that an increase in egg length (likely in addition to the increase in egg width) has contributed to the higher egg volume. In this study, we use egg length as a proxy of egg size which we can measure accurately and with high throughput using the method we have developed.

### Maintenance of *Drosophila* strains

Assays for egg length distribution were performed using 12 inbred lines of *D. simulans* (Signor et al., 2018), Lausanne5 (*D. melanogaster*), Dere01 (*D. erecta*), Dsan01 (*D. santomea*) and Dmau151(*D. mauritiana*). Except for *D. simulans*, all inbred lines were obtained from Bloomington stock center. The rest of assays were performed using an outbred population that was established from 97 *D. simulans* inbred lines (Signor et al., 2018) using a round-robin crossing scheme. We maintained the crosses for 15 generations at 25°C and mixed an equal number of crosses to set up the outbred population. The outbred population and *Drosophila* strains was maintained on standard *Drosophila* medium at 25°C with 12hrs light and 12hrs dark daily and 50-60 % humidity. Larval crowding (Venkitachalam et al., 2022) and temperature (Avelar, 1993; Azevedo et al., 1996; Imai, 1932) affect *D. melanogaster* egg size. To reduce the environmental effects we maintained the fly cultures at low density (400 eggs/bottle) using a pipetting method (Nouhaud et al., 2018) for at least two generations before the egg size measurement. In brief, the volume of washed eggs was compared to a series of tubes containing a known volume of 1x PBS ranging from 50-70 μl. The volume of eggs was adjusted to 60 μl, then 1200 μm 1x PBS, i.e. 20x the volume of eggs, was added. 200 μl of this suspension contained on average 400 eggs which were transferred to a bottle. The protocol can be adjusted to account for the difference in egg volume across different species and populations.

### Preparation of *Drosophila* eggs for flow cytometry

Around 500 4-6 days old individuals (~ 50:50 sex ratio) were transferred to an embryo collection cage (Cat. #59-101, Genesee Scientific) with an 10 mm agar plate (4% agar and 4% sucrose) topped with yeast paste to facilitate oviposition (Becher et al., 2012). Although the age of females doesn’t affect the egg length (Imai, 1932; Lott et al., 2007; Warren, 1924) we restricted the measurements to eggs laid by 4-6 days old females. We limited the oviposition period to 17-19 hours to prevent the hatching of the eggs. A few drops of lukewarm water were added to the agar plate to dissolve the yeast paste, and the solution of yeast and water together with the eggs was transferred to a fine net using a brush. Eggs were washed with water until all yeast grains were dissolved, the eggs were then transferred to a 1.5 ml tube containing 600 μl 1x PBS.

### Size measurement and sorting using large particle flow cytometry

We ran all our samples on a Biosorter® (Union Biometrica) machine with fluidics and optics core assembly units (FOCA) 1000 μm using 1x PBS as a sheath solution. To prevent clogging the flow cell, we re-suspended the eggs in 1x PBS such that the final density of ~400 eggs/ml was reached. We used the pipetting method (Nouhaud et al., 2018) described above to adjust the egg density. Samples were stirred continuously using a mechanical stirrer in a 50 ml sample cup and traveled to the flow cell surrounded by the sheath solution (1x PBS) that focused them into the center of the stream where they were interrogated by multiple lasers. We ran Biosorter® using FlowPilot™ software using a pressure sample cup 0.60-0.65 and pressure diverter 1.2. If samples were to be sorted, a droplet containing fluid and the sorted object was generated which dropped into the collecting container. We sorted samples using the following parameters: drop width of 22 milliseconds (ms) and sort delay of 28 ms. We cleaned the mechanical stirrer with distilled water and the system with 1x PBS after running each sample to avoid cross-contamination.

Large particle flow cytometer measures two optical characteristics of the objects: time of flight (TOF) which is the time each object blocks the light, and extinction (EXT) which is determined by the total integrated signal of the blocked light (Fig. S1). We measured the TOF of polystyrene reference beads (200 and 430 μm) and fitted a linear regression model using TOF as the response variable and size as the explanatory variable. Using the estimated intercept and slope, TOF measurements of all samples were converted to size (size = 5.94 TOF - 491.78). FlowPilot™ software allows real-time visualization of EXT and TOF of objects as scatter plots and regions of interest, i.e. gates, can be chosen on this plot for sorting objects (Fig. S1). Based on the estimated size using TOF measurements, multiple gates were set up that corresponded to different size ranges (Fig. S1). These gates were used for sorting eggs which were dispensed in 1x PBS.

### In silico filtering of debris particles

The size distribution of eggs in all analyzed samples was bimodal (‘No filtering’ curve in Fig. 1C). Manual inspection of the objects in the first mode revealed that the objects are mostly debris and yeast particles while the particles in the second mode were all *Drosophila* eggs. We thus filtered out the objects that corresponded to the first mode by first fitting a kernel density distribution to the size distribution and then identifying the border between the two modes. In brief, we fitted kernel density estimation that is controlled by a smoothing parameter, i.e. bandwidth, to the size distribution. To identify a suitable bandwidth for smoothing the density distribution we performed an exhaustive search over a range of bandwidths by fitting the kernel density function with a Gaussian shape. We performed K-Folds cross-validation by splitting the dataset into train and test sets. We split data into five folds; four folds formed the training set and one fold was used as a validation set. The best bandwidth giving the highest score was chosen as the bandwidth to smooth the kernel density distribution. We then computed the log-likelihood of each dataset under the fitted kernel density distribution and identified the border between the two modes as the first valley in the log-likelihood. The border between the two modes was used as the empirical size threshold for filtering yeast particles for each sample (‘Debris filtering’ curve in Fig. 1C).

### In silico filtering of misaligned eggs

The length of a fraction of eggs was estimated to be 200-400 μm (‘Debris filtering’ curve in Fig. 1C) by LPFC in all the samples which is smaller than the range of *Drosophila* egg length, i.e. 400 to 600 μm (Imai, 1932; Lott et al., 2007; Markow et al., 2009; Venkitachalam et al., 2022; Warren, 1924). Manual measurement of these eggs showed that their size was underestimated by LPFC likely due to the orientation of eggs during measurement. When an prolate ellipsoid object passes through the laser path along its long axis the recorded TOF corresponds to its length (Fig. 1A). But if the object is misaligned, i.e. the object doesn’t pass along the long axis, TOF and subsequently length will be underestimated (Fig. 1B). Using a more viscous sheath solution, e.g. 1x PBS with 4% methyl cellulose, will reduce the flow rate and facilitate the alignment of objects along their long axis when passing through the laser path (Fig. S2). However, the viscosity of the sheath solution dramatically decreases the speed of analysis.

Thus, in addition to the debris filtering, we used the optical density information to filter out misaligned eggs that were not aligned on their long axis. The optical density which measures the extinction of objects varies as a function of the structure of objects (Fig. 1A-B). We extracted the maximum optical density recorded for an object (W) and the total number of extinction measurements along each object’s time of flight (L) and computed the ellipticalness index 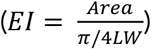 which is the ratio of the area under the optical density profile of an object to the area of the closest elliptical profile (Li et al., 2022; Morales et al., 2020). Due to the elliptical shape of the *Drosophila* egg, an egg aligned on the long axis has a smaller W/L ratio than eggs aligned on the short axis (Fig. 1A-B). For each sample, we computed EI and W/L for all objects and retained objects distributed within half of the standard deviation around the median of EI and W/L. Lastly, after filtering objects based on EI and W/L, the first mode of the distribution of *Drosophila* egg length, which corresponds to the misaligned eggs, was filtered out similarly to the debris filtering step described above.

### Manual measurement of egg length

To validate the accuracy of the estimated egg length using LPFC, we measured the length of eggs manually and automatically using LPFC. Around 2000 individuals from the outbred *D. simulans* population were transferred to an embryo collection cage with 10 mm agar plates topped with yeast paste and laid eggs for 17-19 hours. We prepared eggs for flow cytometry as described above, and sorted 74 eggs from 6 gates corresponding to 400-600 μm (with intervals of 50 μm) in a 96-well plate. During the in-silico filtering (described above) the misaligned eggs with the estimated average length of 362 μm (N = 16 samples, Table S2) by Biosorter® were filtered out. Therefore, sorting eggs from gates with the estimated length of 400-600 μm assures the absence of misaligned eggs. We then manually measured the length of the sorted eggs. We transferred the eggs to a 100 mm petri-dish containing a dark medium (4% agar and 0.5% charcoal), positioned the eggs on their dorsal side, and took high-resolution photos using a Leica stereomicroscope (M205 FA). We measured the egg length manually using ImageJ (Schneider et al., 2012). The scale of images was transformed to μm and the egg length was measured by drawing a line along the longitudinal axis of each egg. The concordance between the manual length measurements and the automated length estimates by LPFC was tested using Spearman’s rank correlation.

Moreover, we tested whether submersion in PBS influences the egg length measurement. We positioned 46 eggs on their dorsal side on a 100 mm petri-dish containing a dark medium and took high-resolution photos. We submerged the eggs in a 96-well plate for 2.5 hours, then transferred the eggs to a dark medium and took high-resolution photos. We measured the egg length manually using ImageJ as described above. The concordance between the egg length measurements before and after submersion in PBS was tested using Pearson’s correlation.

### Egg-to-adult viability assay

During the egg length measurement and sorting using LPFC eggs might be immersed in PBS for up to several hours. Moreover, while LPFC employs a slower flow rate and lower pressure than traditional flow cytometry to ensure that the sorted objects are viable, passing through the flow cytometer may affect the survival of the eggs. We performed an egg-to-adult viability assay to determine the effect of submersion in PBS and passing through the flow cell on the development of eggs into adults. We collected eggs laid by an outbred *D. simulans* population for 17-19 hours, as described above in the Manual measurement of egg length. To determine the egg-to-adult viability of the eggs without exposure to PBS, the eggs on the agar plate were not washed with PBS (‘no PBS’ treatment); 5 independent sets of 50 eggs were manually counted and transferred to 5 vials with a brush. We then washed the eggs laid on the yeast paste and transferred around 30 μl eggs to three 1.5 ml tubes containing 600 μl 1x PBS; 2 tubes corresponded to ‘0-hour’ and ‘4-hour’ treatments and 1 tube to ‘sorted 4-hour’ treatment. The eggs in 0-hour and 4-hour treatments were maintained at room temperature (25 °C) for only 20 minutes and 4 hours, respectively. For each treatment, 5 independent sets of 50 eggs were manually counted under a stereoscope and each set was transferred to a vial containing standard *Drosophila* medium. We ran the eggs in the ‘sorted 4-hour’ treatment on LPFC and sorted 7 independent sets of 50 eggs from gates corresponding to 400-600 μm, i.e. the whole range of egg length distribution. Each set was transferred to a vial. The number of eclosed adults from each vial for all the treatments was counted on 4 consecutive days until no more flies were eclosed.

To test for the effect of PBS submersion and sorting by LPFC on the egg-to-adult viability we fitted a generalized linear mixed model (using a binomial distribution) with treatment, i.e. submersion time and sort status, as a fixed categorical effect with 4 levels (no PBS, 0-hour, 4-hour, and sorted 4-hour). The used model is as follows: success_ij_ ~ μ + treatment_i_ + error_ij_, where success is a matrix with the number of survived and dead eggs. All treatments have a sample size of 5 replicates except for the sorted 4-hour treatment which has 7 replicates. The replicates were included as a random effect. The significance of the fixed effect was tested using ANOVA F-tests and we used Tukey’s HSD to correct for multiple testing.

## Results

### Egg length distribution after in-silico filtering of debris particles and misaligned eggs

The length distribution of all samples was bimodal (Fig. 1C) and the average size of the 1st and 2^nd^ modes were 92.38 and 412.21 μm (n = 16 samples, Table S1). The objects in the first mode were mostly debris and yeast particles upon manual inspection. We used the computed border between the two modes as an empirical size threshold to remove the objects in the 1^st^ mode. These sample-specific empirical thresholds ranged from 92.9 to 212.21 μm with an average of 184.29 μm (n = 16 samples, Table S1).

The 2^nd^ mode with an average size of 412.21 μm (n = 16 samples) corresponds to the length of *Drosophila* eggs (Imai, 1932; Lott et al., 2007; Markow et al., 2009; Venkitachalam et al., 2022; Warren, 1924) but also contained some misaligned eggs (Fig. 1A-B). The length of misaligned eggs was estimated between 200 and 400 μm by LPFC (first peak in the ‘Debris filtering’ curve in Fig. 1C) but was underestimated likely due to the improper orientation of eggs during measurement. To filter the misaligned eggs, we computed the ellipticalness index (EI) and the ratio of the maximum optical density of an object to the number of extinction measurements along an object’s time of flight (W/L) values using the optical density data for each sample. Each dataset was further filtered by removing eggs that were not aligned on the long axis (Fig. 1B) by retaining objects with EI and W/L values within half of the standard deviation around the median. As the final step for filtering the misaligned eggs, we removed the objects corresponding to the 1^st^ peak of the egg distribution (first peak in ‘Debris filtering’ curve in Fig. 1C). These filtering steps did not remove objects with the large length showing that EI and W/L parameters properly identify misaligned eggs where the length in underestimated (Fig. 1C). The computed sample-specific empirical threshold for filtering these objects ranged from 163.36 to 317.12 μm, with an average of 317.12 μm (n = 16 samples, Table S2).

### Accuracy and throughput of egg length measurement and sorting

We validated the accuracy of egg length measurements and sorting using LPFC by comparing the estimated egg length by LPFC with the manual measurements. We sorted eggs from several gates corresponding to 400 to 600 μm, where misaligned eggs are absent, and measured the length of the eggs manually. We observed a high correlation between the manual measurements and the estimated length using LPFC (Fig. 2A). We further assessed the effect of submersion in PBS on the egg length. The manually measured egg length before and after 2.5 hours of submersion in PBS was highly correlated (Fig. 2B).

**Fig. 2.**
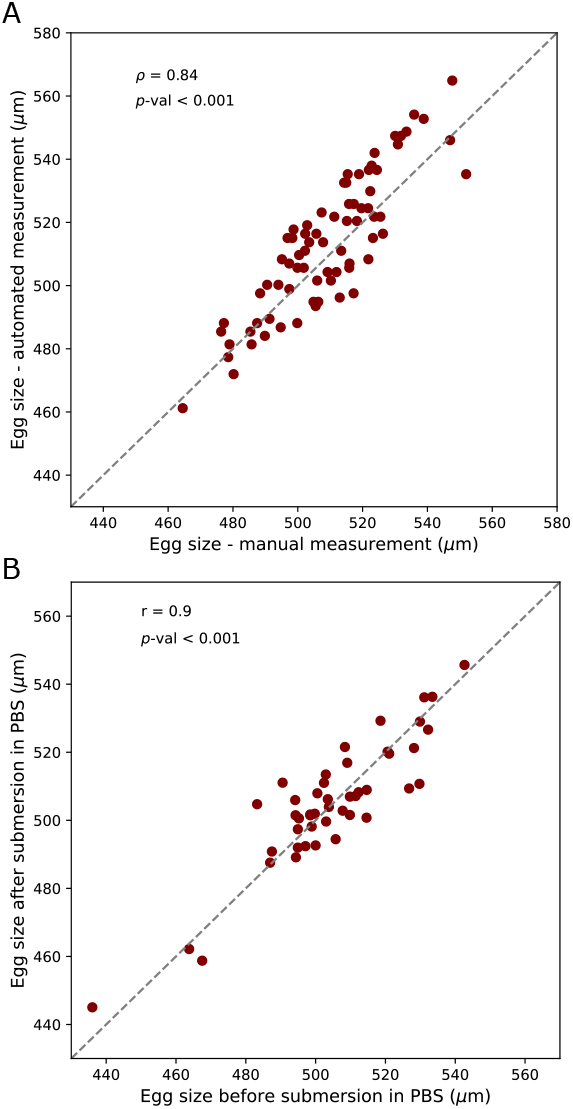
Accuracy of the estimated egg length using large particle flow cytometry. A) The estimated sizes through manual measurement and large particle flow cytometry are highly correlated (Spearman’s rank correlation **ρ** = 0.84, *p*-value < 0.001). B) Submersion in PBS does not affect the egg length as the measured egg length is highly correlated before and after submersion in PBS (Pearson’s correlation r = 0.9, *p*-value < 0.001). The slope of the dashed lines in A and B is 1.

The measurement of egg length is high throughput and fast. We measured an average of 316 eggs/min which corresponds to 214 eggs/min (n = 13 samples) after filtering out the misaligned eggs (Fig. 3A). We evaluated the efficiency of sorting as sort recovery which is the percentage of objects that were sorted and dispensed as a fraction of all the objects fulfilling sorting criteria, i.e. size. In our hands, the median sort recovery was 63.54% (n = 28 samples, Fig. 3B) and the median speed of sorting was 69 eggs/min (n = 28 samples, Fig. 3C).

**Fig. 3.**
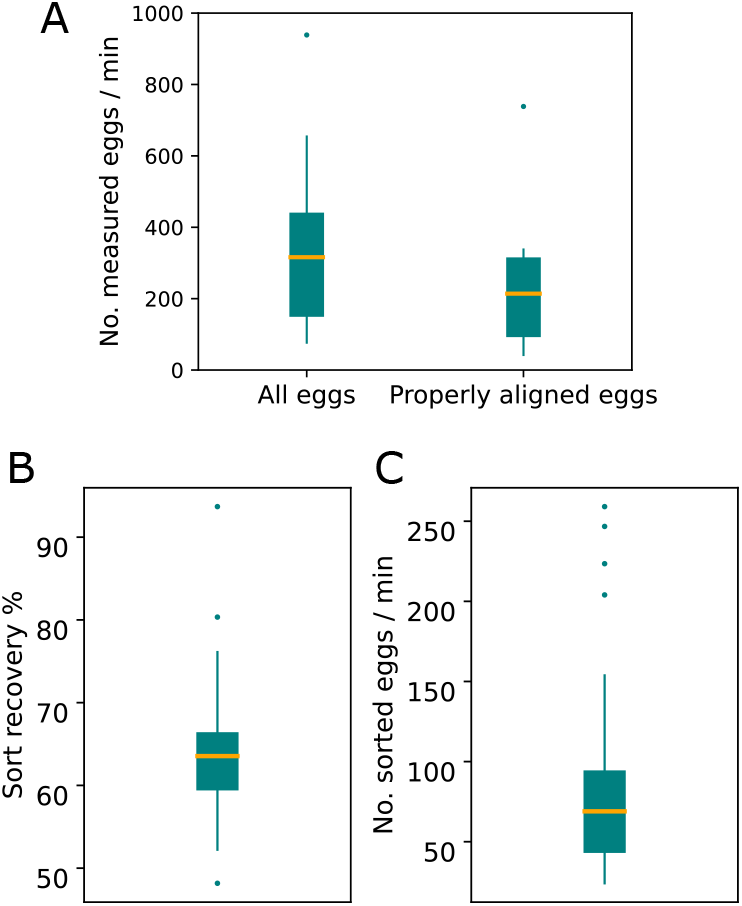
Throughput of the size measurement and sorting of eggs using large particle flow cytometry. A) The median speed of egg size measurement is 316 eggs/min after filtering for debris (‘Debris filtering’ curve in Fig. 1C) and 214 eggs/min (n = 13 samples) after filtering the misaligned eggs (‘Misaligned filtering’ curve in Fig. 1C). B) The median sort recovery as an estimate of the efficiency of sorting is 63.54% (n = 28 samples). C) The median speed of sorting is 69 eggs/mins (n = 28 samples).

### The effect of handling and sorting by large particle flow cytometer on egg viability

To demonstrate the suitability of using LPFC for the measurement and sorting of viable eggs, we tested whether the sorted eggs develop to adults normally. We observed no significant difference in egg-to-adult viability between eggs that were not washed with PBS (42.4±3.36 out of 50) and those that were submerged in PBS briefly for 20 minutes (40.74±6.08 out of 50) and 4 hours (43.2±5.12 out of 50) (Fig. 4, a generalized linear mixed model followed by Tukey’s HSD test with correction for multiple testing, *P*_no-PBS-0hour_ = 0.7307, *P*_no-PBS-4hour_ = 0.9567, *P*_0hour-4hour_ = 0.4405). Moreover, eggs that passed through LPFC showed no decrease in egg-to-adult viability (42.57±3.6 out of 50) compared to the eggs submerged in 1x PBS for the same duration of time (43.2±5.12 out of 50) (Fig. 4, *P*_no-PBS-sorted 4hour_ = 1.0000, *P*_*0hour*-sorted 4hour_ = 0.6673, *P*_4hour-sorted 4hour_ = 0.9540).

**Fig. 4.**
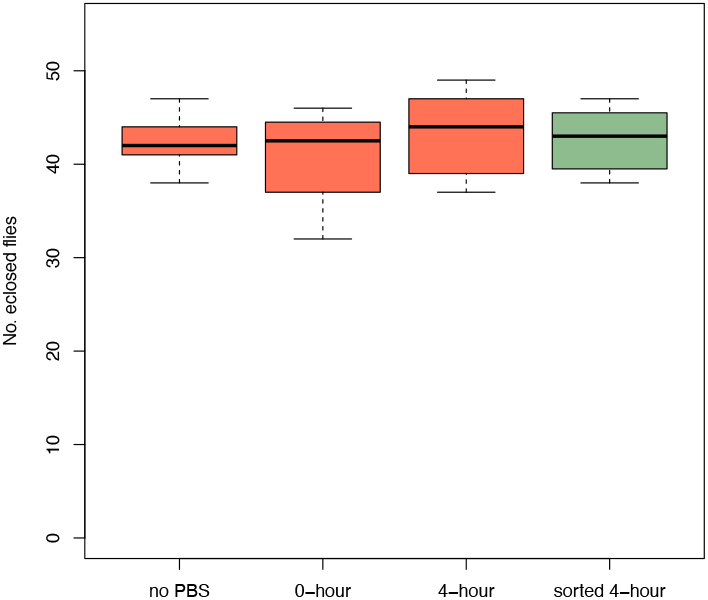
The viability of eggs is not reduced by submersion in PBS and sorting by large particle flow cytometry. There is no significant difference in the viability of eggs that were not washed with PBS (no PBS), with those that were submerged in PBS for 20 minutes (0-hour), and 4 hours (4-hour). Passing through the flow cytometer (sorted 4-hour) does not reduce the egg-to-adult viability either (a Generalized linear mixed model followed by Tukey’s HSD test with correction for multiple testing, *P*_no-PBS-0hour_ = 0.7307, *P*_no-PBS-4hour_ = 0.9567, *P*_0hour-4hour_ = 0.4405, *P*_no-PBS-sorted 4hour_ = 1.0000, *P*_*0hour*-sorted 4hour_ = 0.6673, *P*_4hour-sorted 4hour_ = 0.9540).

## Discussion

Here, we establish a method for accurate and high throughput measurement of *Drosophila* egg length using large particle flow cytometry. The measurement of egg length is highly accurate (Fig. 2), fast and high throughput (Fig. 3A). Viable eggs within a specified length range can be sorted rapidly (Fig. 3C) with no reduction in egg-to-adult viability (Fig. 4).

### Applicability of measuring *Drosophila* egg length using LPFC

Obtaining accurate and high throughput size measurements of *Drosophila* eggs has several applications. First, this high throughput method is a suitable approach for the use of *Drosophila* egg length as a life history trait in large-scale studies where many populations should be analyzed in parallel. Second, this approach allows high throughput analysis of the variation in egg length in natural populations of different species of *Drosophila*. Third, the high throughput of egg length measurement using large particle flow cytometry coupled with the available reference panels of *D. melanogaster* (Mackay et al., 2012) and *D. simulans* (Signor et al., 2018) provide an opportunity to investigate the genetic architecture of egg length variation through QTL mapping and GWAS. Fourth, this method provides the possibility of sorting viable *Drosophila* eggs from the range of the length distribution and performing selection experiments with large populations size and replicates.

### Recommendations for the use of LPFC for size measurement

The opportunities provided by LPFC for the measurement of the organism’s size are not limited to *Drosophila* eggs and can be applied to any organism within the detectable range of this instrument (10-1500 μm). Our recommendation is to perform preliminary experiments to optimize the settings of the instrument for the specific sample and application. Some of the important settings for the optimization of the measurement and sorting include flow rate, mixing speed, and sample density. The flow rate is adjusted by the pressure of the sample cup and determines the number of objects analyzed in a unit of time. The Biosorter® user manual recommends adjusting the flow rate to achieve 10 to 30 objects per second. The speed of mixing should be adjusted to assure objects are not damaged but are also homogenously suspended in the buffer. The appropriate sample density is a balance between the speed and accuracy of measurement and sorting. Highly dense samples can potentially clog the flow cell and also result in inaccurate size measurement if objects cannot be measured separately by the instrument. On the one hand, contaminated events, i.e. the objects which don’t meet the sort criteria are with the sortable object in the same drop, cannot be sorted. Thus, too high density may also reduce the sort recovery. On the other hand, highly diluted samples will make the analysis lengthy. The weight of objects affects the flow rate and mixing speed of the analysis. For objects that are dense and heavy such as *Drosophila* eggs, the mixing speed should be fast to keep the objects suspended. Moreover, the pressure of the sample cup which affects the flow rate cannot be too low as objects may not even enter the flow cell if too low pressure is applied.

The number of eggs in each run should be adjusted to assure a reliable estimation of the size distribution. Our in-silico filtering algorithm has identified distinct fractions corresponding to debris and misaligned eggs for samples containing 1000 to 16,000 eggs (Table S2). However, the filtering algorithm may require additional optimization if too few objects are available.

The settings of the instrument and the filtering steps should be adjusted according to the size and shape of objects. If the object’s size is too small that overlaps with the size of debris, more intensive cleaning for removing debris may be required. For objects that have an approximately spheroid shape such as *Arabidopsis* seeds (Cervantes et al., 2010) the orientation of the object does not affect the estimated size by LPFC (Morales et al., 2020). However, different instrument settings and in silico filtering steps are needed for the objects with shapes that deviate from a sphere. The method developed in this study is optimized for measuring the length (the axis of rotational symmetry) of prolate ellipsoid objects (such as *D. melanogaster* species subgroup) or the width (the axis perpendicular to length) of oblate ellipsoid objects. We recommend a similar in-silico filtering step used in this study to remove debris and misaligned objects. Additionally, using a viscous sheath solution to reduce the flow rate will minimize the number of misaligned objects (Fig. S2). However, a lower flow rate will reduce the speed of analysis. Moreover, the average maximum height of optical density (W), which we used for filtering misaligned objects, can also be used in combination with other sorting criteria, e.g. size, to increase the efficiency of sorting objects. If measuring the width of a prolate ellipsoid object or the length of an oblate ellipsoid object is needed, the run specifications should be optimized to maximize the fraction of objects in the desired orientation to be measured by LPFC. Moreover, the in-silico filtering steps should be adjusted to remove the misaligned objects. We recommend the sorting of objects according to the size be performed from the size range (i.e. gate) that the absence of misaligned objects is assured.

### Caveats

Large particle flow cytometers are expensive instruments that may not be easily acquired by small research groups. However, many research facilities house such instruments that can provide access to researchers. Most research facilities have trained personnel that can assist and train researchers.

## Author Contributions

Neda Barghi conceived the ideas and designed the methodology; Neda Barghi and Claudia Ramirez-Lanzas collected the data; Neda Barghi analyzed the data and led the writing of the manuscript. All authors contributed critically to the drafts and gave final approval for publication.

## Data Availability Statement

All the data and code to reproduce the results are deposited in https://github.com/NedaBarghi/EggSizeBiosorterProtocol.

## Acknowledgment

The BioSorter® Large Particle Flow Cytometer (Union Biometrica, Holliston, MA) of the Max Perutz Labs BioOptics FACS Facility (https://www.maxperutzlabs.ac.at/research/facilities/biooptics-facs) was used for sorting. We thank Johanna Stranner and Kitti Dora Csalyi for their help in optimizing the sorting protocol. We thank Paula Marconi for her help in the egg-to-adult viability assay. We thank Emmanouil Lirakis for assistance with using the Leica stereomicroscope. This work was supported by the Austrian Science Fund (FWF, P 32672) to NB. We thank Sara Signor (North Dakota State University) for providing *D. simulans* inbred lines. *Drosophila* strains Lausanne5 (*D. melanogaster*), Dere01 (*D. erecta*), Dsan01 (*D. santomea*) and Dmau151 (*D. mauritiana*) are obtained from Bloomington stock center by Robert Kofler. We thank Samuel Church and the anonymous reviewer(s) for constructive comments on the manuscript.

## Supplementary figures and tables

**Fig. S1.**
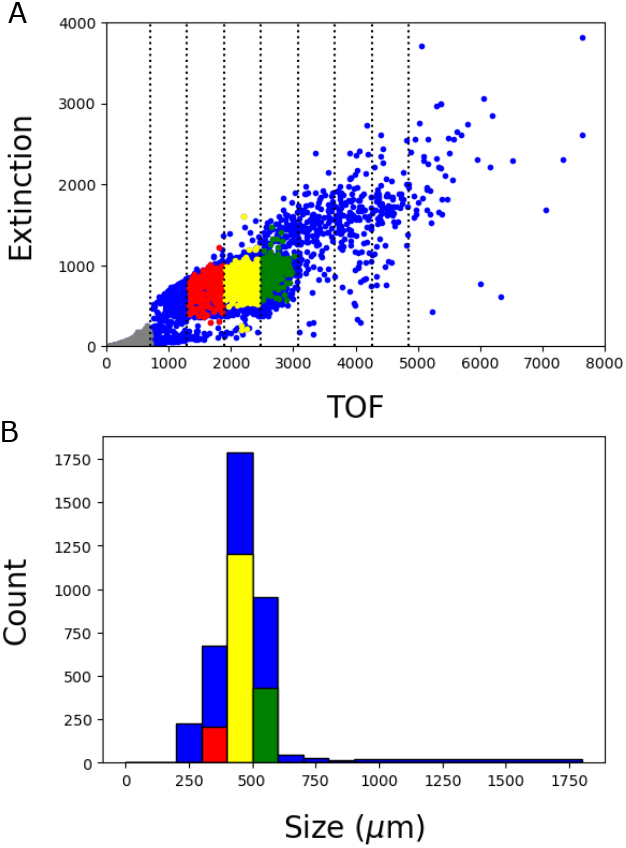
Time of flight (TOF), the measured time an object blocks the light, is used to estimate the object’s size. *Drosophila* eggs are washed and dispensed in 1x PBS buffer and run on large particle flow cytometry. TOF and extinction are measured (A) and TOF is converted to size (μm) using the TOF data of reference beads. Based on the size distribution, user-defined sorting gates are set up (depicted as dotted lines in (A), these gates correspond to size 200-900 μm with intervals of 100μm), and the required number of eggs from each gate can be sorted. Red, yellow and green bars in (B) show the number of sorted eggs in these 3 size gates in (A).

**Fig. S2.**
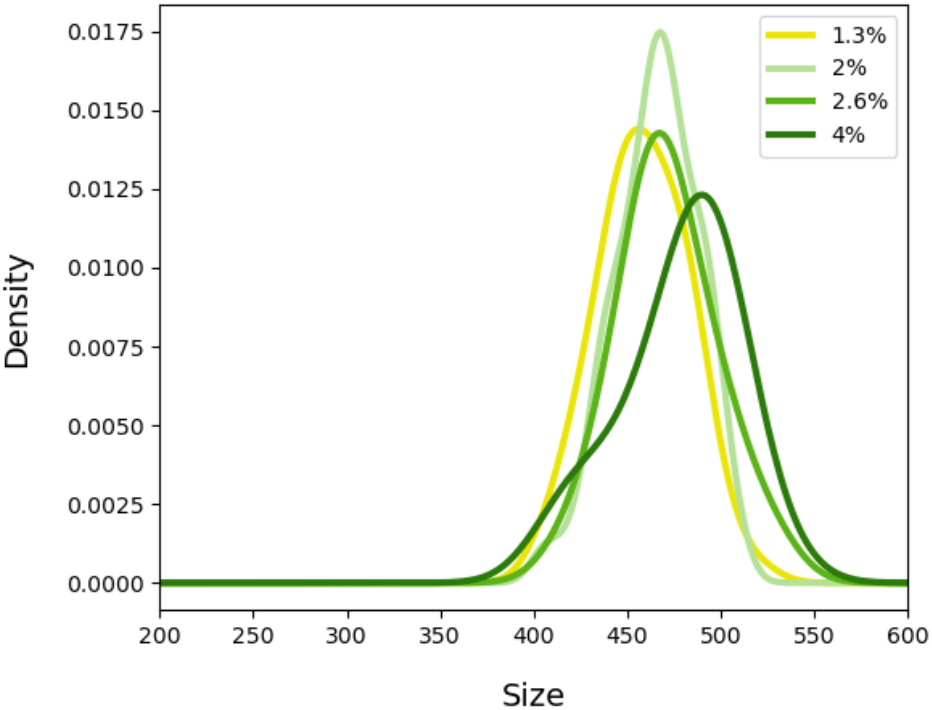
Distribution of the egg length estimated using different concentrations of methyl cellulose (MC) in 1x PBS as sheath buffer. As the concentration of MC increases the sheath buffer becomes more viscous which will reduce the flow rate. The slower flow rate facilitates the alignment of objects along the long axis while passing through the laser path. However, the increased viscosity of the sheath buffer dramatically reduces the speed of analysis. The same sample has been used for all runs shown in the plot.

**Fig. S3.**
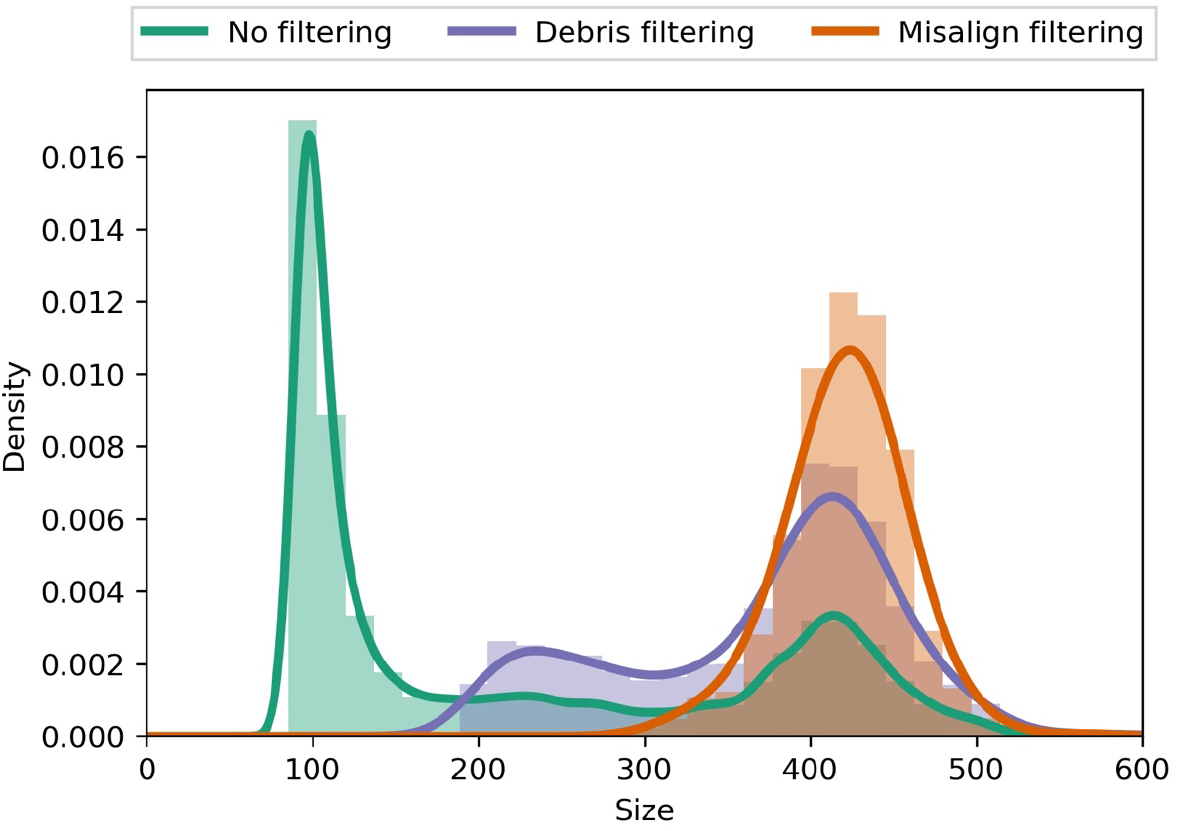
Distribution of the egg length in sample Dsim196 before and after in silico filtering of debris and misaligned objects. The initial size distribution is bimodal (No filtering). The 1^st^ mode corresponds to debris and yeast particles, and ‘Debris filtering’ remove these small objects. Further filtering steps based on EI and W/L parameters (Misalign filtering) removed the misaligned eggs. Size is in μm units. The statistics of egg length distribution is presented in Table S1 and S2.

**Table S1.** The statistics of egg length distribution. Size threshold is used to remove 1^st^ peak containing egg debris from the 2^nd^ mode in the first filtering step. Dsim#, #: stock number (*D. simulans*), Lausanne5 (*D. melanogaster*), Dere01 (D. *erecta*), Dsan01 (D. *santomea*) and Dmau151 (*D. mauritiana*). Except for *D. simulans*, other inbred lines are obtained from Bloomington stock center. *D. simulans* inbred lines are provded by Sara Signor (North Dakota State University).

| Sample | 1st peak mode (μm) | size threshold (μm) | 2nd peak mode (μm) |
| --- | --- | --- | --- |
| Dsim1 | 94.903 | 212.212 | 449.078 |
| Dsim4 | 92.209 | 176.977 | 459.851 |
| Dsim5 | 92.209 | 196.196 | 427.531 |
| Dsim11 | 92.209 | 177.778 | 426.184 |
| Dsim75 | 92.209 | 207.407 | 450.425 |
| Dsim90 | 92.209 | 173.774 | 427.531 |
| Dsim91 | 92.209 | 92.893 | 93.556 |
| Dsim146 | 92.209 | 180.981 | 450.425 |
| Dsim166 | 92.209 | 198.599 | 436.958 |
| Dsim185 | 92.209 | 201.802 | 426.184 |
| Dsim196 | 92.209 | 194.595 | 423.491 |
| Dsim237 | 92.209 | 182.583 | 431.571 |
| Lausanne5 | 92.209 | 146.547 | 420.798 |
| Dere01 | 92.209 | 207.407 | 431.571 |
| Dsan01 | 92.209 | 211.411 | 384.438 |
| Dmau151 | 92.209 | 187.387 | 455.811 |

**Table S2.**
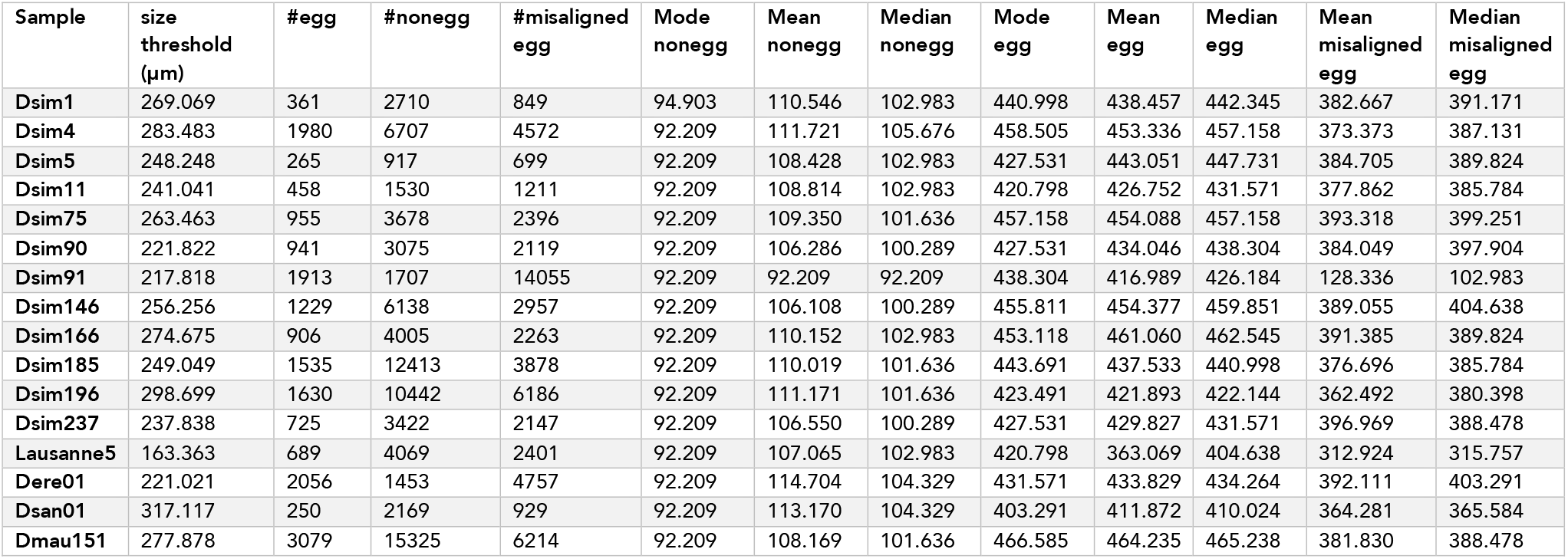
The statistics of egg length distribution filtered using sample-specific threshold. Each dataset is filtered as described in Fig. 1. The 1^st^ filtering step is based on the size threshold in Table S1. The filtered objects at this step are mostly egg debris. # nonegg: the number of small objects filtered in the first filtering stage. The mode, mean and median of these objects are Mode nonegg, Mean nonegg, and Median nonegg respectively. The second filtering step involved removing the misaligned eggs. The number, mean and median of these misaligned eggs are shown as #misaligned egg, Mean misaligned egg and Median misaligned egg. The last filtering step is to remove the objects in the first peak of distribution after removal of the misaligned eggs. Size threshold (μm) is used for this filtering step. #egg, Mode egg, Mean egg and Median egg are the number, mode, mean and median of the eggs in the final dataset. Dsim#, #: stock number (*D. simulans*), Lausanne5 (*D. melanogaster*), Dere01 (*D. erecta*), Dsan01 (*D. santomea*) and Dmau151 (*D. mauritiana*). Except for *D. simulans*, other inbred lines are obtained from Bloomington stock center. *D. simulans* inbred lines are provded by Sara Signor (North Dakota State University).

